# Metabolic activity affects response of single cells to a nutrient switch in structured populations

**DOI:** 10.1101/580563

**Authors:** Alma Dal Co, Martin Ackermann, Simon van Vliet

## Abstract

Microbes live in ever-changing environments where they need to adapt their metabolism to different nutrient conditions. Many studies have characterized the response of genetically identical cells to nutrient switches in homogenous cultures, however in nature microbes often live in spatially structured groups such as biofilms where cells can create metabolic gradients by consuming and releasing nutrients. Consequently, cells experience different local microenvironments and vary in their phenotype. How does this phenotypic variation affect the ability of cells to cope with nutrient switches? Here we address this question by growing dense populations of Escherichia coli in microfluidic chambers and studying a switch from glucose to acetate at the single cell level. Before the switch, cells vary in their metabolic activity: some grow on glucose while others cross-feed on acetate. After the switch, only few cells can resume growth after a period of lag. The probability to resume growth depends on a cells’ phenotype prior to the switch: it is highest for cells crossfeeding on acetate, while it depends in a non-monotonic way on growth rate for cells growing on glucose. Our results suggest that the strong phenotypic variation in spatially structured populations might enhance their ability to cope with fluctuating environments.

## Introduction

Environmental conditions are constantly changing on earth and a central prerequisite for any lifeform is the ability to cope with these fluctuations. Microorganisms living in e.g. the soil or animal gut experience strong fluctuations in the quantity and quality of nutrients in their environment. To deal with these fluctuations, microorganisms have evolved a large variety of mechanisms for adapting their metabolism to new nutrient conditions [1].

Recent studies have shown that genetically identical cells can vary in the time they need to adapt to the same nutrient switch (lag time), and often a fraction of cells cannot adapt at all [2–9]. This heterogeneity in response is primarily due to phenotypic differences among cells at the time of the switch, for instance in their protein levels or metabolic fluxes [2–10]. For example, the ability of *Escherichia coli* cells to switch from growth on glucose to growth on lactose depends on how many proteins of the lactose pathway the cells have at the time of the switch [9]. Similarly, their ability to switch from growth on glucose to growth on a gluconeogenic carbon source (e.g. acetate) depends on their gluconeogenic flux at the time of the switch [2]. Whenever cells share a homogeneous environment, differences in protein levels and metabolic fluxes are mainly due to stochastic fluctuations in their gene expression [11,12].

Homogeneous environments are likely a special case in nature, because many bacteria live in dense spatially structured groups, such as biofilms and microcolonies [13]. The high densities in these groups (up to a thousand times higher than in batch cultures) allow cells to modify their local microenvironment with the secretion and uptake of compounds [14,15]. Consequently, cells at different locations adapt to different microenvironments and vary in their gene expression and metabolic fluxes [1,16–23]. In spatially structured populations cells thus differ in phenotype both due to stochastic fluctuations in gene expression (as in homogenous cultures) *and* due to physiological adaptation to different local microenvironments. As a result, structured populations have more phenotypic variation than homogenous populations.

Phenotypic variation can have important consequences for the ability of a population to adapt to environmental changes [24]. Adapting to new environmental conditions can be time consuming or even impossible for cells due to energy or resource limitations [25,26]. If cells differ in their phenotypes, either due to stochastic gene expression or adaptation to local microenvironments, some of them might have an increased ability to grow in the new conditions. As a result, the population as a whole can resume growth faster after the environment changes unexpectedly [12,24,26,27]. The higher degree of phenotypic variation in structured populations might thus have important consequences for the ability of these populations to cope with fluctuating environments.

While much is known about the response of cells to nutrient changes in homogeneous environments like batch cultures, little is known about their response in structured populations. How quickly do bacteria inside a structured population respond to changes in the environment? How heterogenous is their response? Here, we addressed these questions by growing dense populations of *E. coli* in microfluidic devices and studying how they cope with a nutrient switch from glucose to acetate.

## Results and Discussion

### Subpopulations specialize on different metabolic activities in spatially structured populations

We grew *E. coli* cells inside microfluidic chambers that host about 1000 individuals in closely packed two-dimensional populations (figure 1a). The chambers open on one side into a flow channel where nutrients are supplied (figure 1a). This setup allows us to switch nutrients, in our case from glucose to acetate, in a tightly controlled manner. We characterized the phenotype of each cell inside the population by measuring their growth rate, and the expression of two metabolic genes: *ptsG*, a high affinity glucose importer expressed when glucose concentrations are low [28–30], and *acs*, a gene involved in acetate metabolism expressed when catabolite repression (i.e. inhibition of catabolism of carbon sources other than glucose) is released [29,31,32].

**Figure 1.**
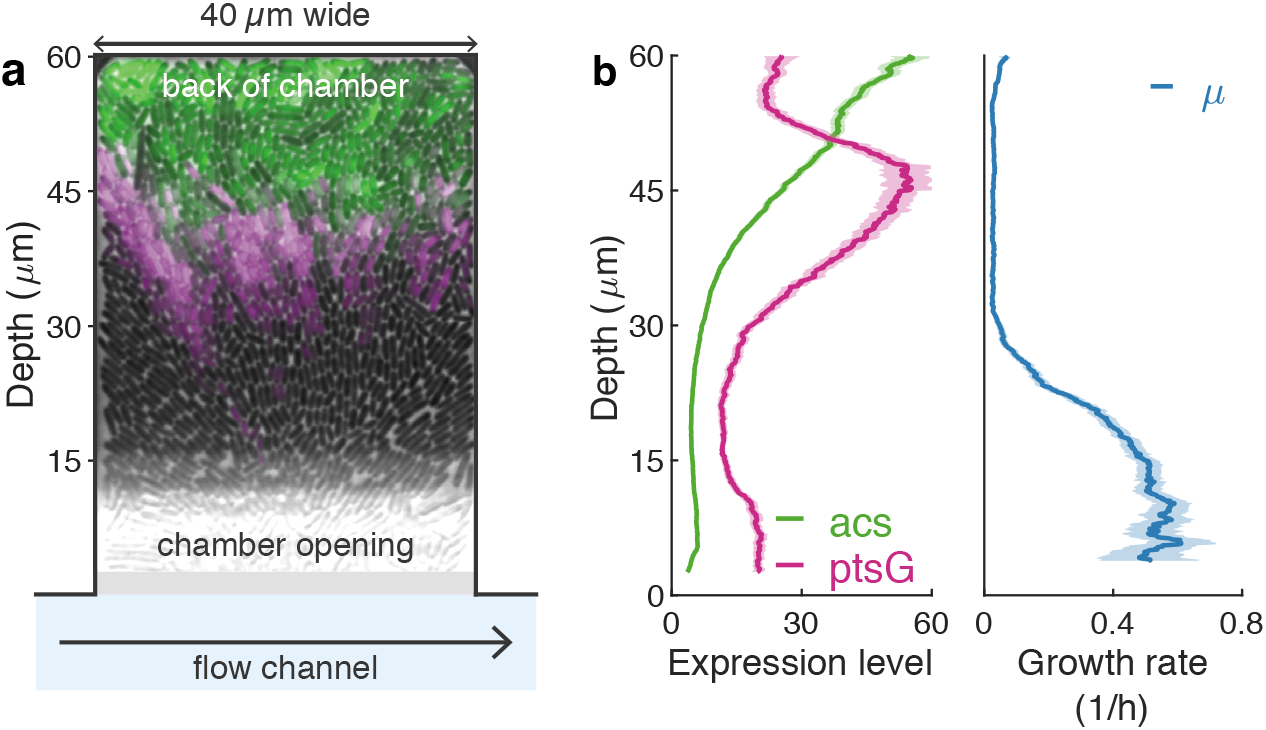
Emergent nutrient gradients create phenotypic variation. **(a)** Cells form closely packed mono-layers of about 1000 cells in microfluidic chambers that open on one side into a flow channel. When a low concentration (800μM) of glucose is supplied, cells create metabolic gradients as they consume glucose and release metabolites. As a result, growth rates and gene expression change with depth. False color image shows in magenta expression level of *ptsG* (expressed when glucose becomes limiting), and in green *acs* (expressed when catabolite repression is released) superimposed on the phase contrast image. **(b)** Cells vary in their expression of *ptsG* (magenta), *acs* (green), and growth rate (blue). For each chamber we calculated the average gene expression and growth rate (both measured at single cell level) as function of depth. Lines show the mean value and shaded areas the 95% confidence interval over 15 chambers. Single cell measurements were averaged over the chamber width and over a moving window with a depth of 3μm.

Gene expression was measures using plasmid based transcriptional reporters, as this allows for nondestructive single cell measurements within a microfluidic setup. However, such reporters have potentially three limitations. First, plasmid copy number variation could lead to overestimate the variation in gene expression between cells. However, previous work has shown that for most promoters copy number variation of the reporter plasmids we used is negligible compared to variation in transcriptional activity [33,34]. Second, because (fluorescent) proteins have long lifetimes, fluorescent intensities in cells do not only depend on current, but also on past, transcriptional activity. Third, transcriptional reporters cannot give any insight into post translational regulation that plays an important role in regulating metabolic activity [17]. As a result, transcriptional reporters cannot give direct information about a cell’s current metabolic fluxes. However, they give information about a cell’s metabolic potential, i.e. about the pathways that the cell expresses. This allows to identify subpopulations of cells that have specialized on different metabolic tasks [29,35].

In previous work, we showed that when glucose is supplied at low concentration in the flow channel (800μM), the combined metabolic activity of the cells creates strong nutrient gradients along the depth of the chamber [36]. The local microenvironment varies over a length scale of a few cell-lengths, and as a result the average phenotype of the cells changes with the depth in the chamber. Cells form distinct subpopulations that specialize on different metabolic pathways: close to the chamber opening, where glucose is abundant, cells partly ferment glucose to acetate [37,38]; deeper in the chamber, where glucose becomes growth-limiting, cells grow progressively slower and start to express *ptsG* (figure 1ab); at the very back of the chamber, where glucose is nearly depleted, cells are released from catabolite repression and express *acs*, which allows them to grow on the acetate excreted by cell at the front (figure 1ab). Although we only measured variation in growth rate and *acs* and *ptsG* expression, cells likely varied in many other aspects of their phenotype.

In this study, we investigate how these structured populations respond to a nutrient switch from glucose to acetate. We hypothesized that cells consuming acetate before the switch would be able to continue growing without any lag after the switch, while cells consuming glucose before the switch would vary in their lag time depending on their pre-switch phenotype. To test these hypotheses, we fist let cells stably form the metabolic gradients by growing them on a medium with a low concentration of glucose (800μM). Subsequently, we switched to a medium with a high concentration of acetate (30mM) as the sole carbon source. We followed the population for 38 hours after the switch and quantified the response of the single cells.

### Only specific subpopulations can resume growth after gluconeogenic switch

Out of a total of about 14’000 cells (in 15 chambers), only 237 cells (1.7%) could grow after a switch to acetate (figure 2a, SI movies 1-4). For these 237 cells, the time required to resume growth (lag time) varied substantially. A total of 69 cells could continue to grow without any lag and most of these cells (63) clustered in the back of the chamber, where they were likely already adapted to consuming acetate (figure 2ab). The remaining 168 cells had a broad distribution of lag times with a median of 4.1 hours (2.4-5.8h inter quartile range), though the lag time could be as long as 25.5 hours (figure 2a).

**Figure 2.**
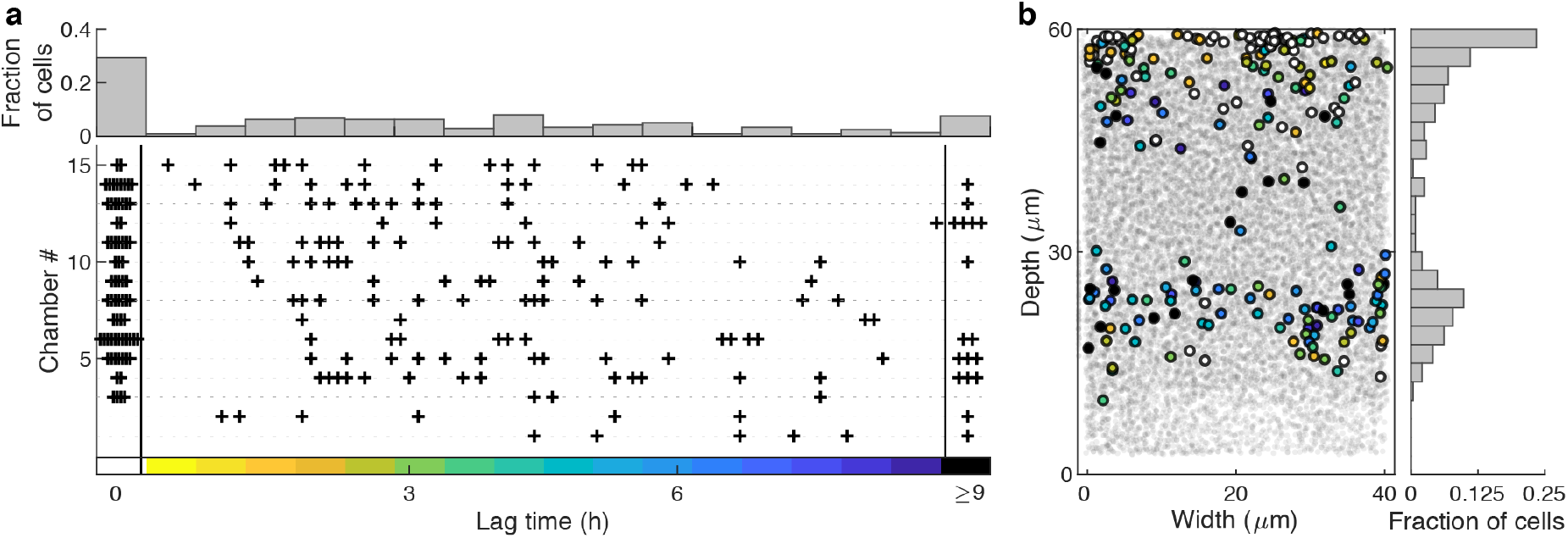
Cell that can grow after switch to acetate reside at specific depths. **(a)** Cells that can grow post-switch vary strongly in their lag time after a switch from the glucose gradient to high amounts of acetate. Data of all 15 chambers (ca. 14’000 cells) was pooled together. **Top:** histogram of lag times; 29% of cells can continue growing without interruption, the other cells lag for 0.3 to 25.5h. **Bottom:** each horizontal line represents a single chamber and each cross indicates the time when a cell resumes growth. Points for lag times of 0h and >9h have a horizontal offset to minimize overlap. **(b)** Cells that can grow post-switch are located either in the back of the chamber or at a depth of 15–30μm. **Left:** the location of all cells that can grow post-switch is marked with large dots and the color indicates their lag time. Small gray dots show the location of all other cells (all 15 chambers shown together). **Right:** histogram of the depth at which cells that can grow post-switch.

Cells that grew after the switch were located either at very back of the chamber or between a depth of around 15 to 30μm (figure 2b). Using a cluster analysis, we found that we could accurately separate these two groups of cells using a threshold depth of 38.5μm (figure 3a). We analyzed the phenotypes of cells that could grow post-switch for both of these clusters (back and front of chamber) separately. Cells that could grow postswitch in the back of the chamber expressed *acs* and grew slowly on acetate before the switch, while cells that could grow post-switch in the front of the chamber did not express *acs* and *ptsG* and grew at high, but non-maximal, rates on glucose (0.4 < *μ* < 0.87h^−1^, figure 3b). All other cells were generally unable to grow after the switch (figure 3b).

**Figure 3.**
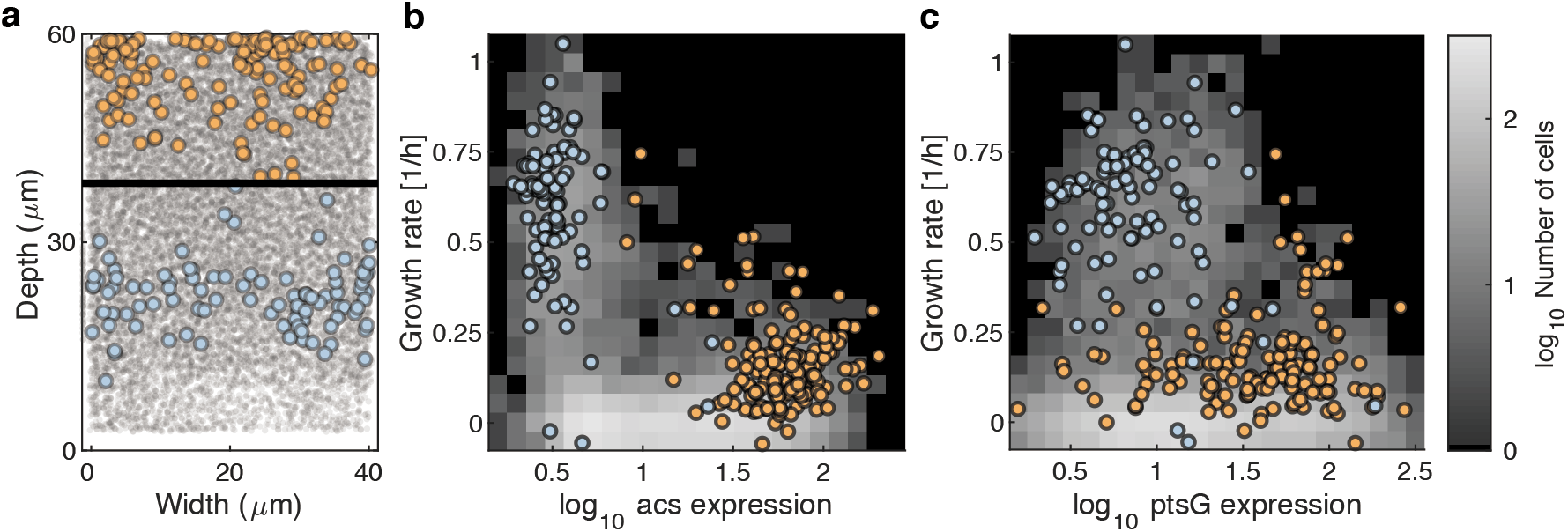
Cell that can grow post-switch come from distinct phenotypic subpopulations. **(a)** Cells that could grow post-switch were classified based on their location in the chamber into two groups, one in the back (depth < 38.5μm, orange) and one in the front of the chamber (blue), using k-means clustering. The data of all 15 chambers is shown together. **(b,c)** The two groups have distinct phenotypes: the subpopulation in the front (purple) is characterized by high, but non-maximal, growth rates and low *acs* expression; the subpopulation in the back (orange) is characterized by intermediate growth rates and high *acs* expression. *pstG* expression is generally higher for cells in the back, but there is considerable overlap between the two groups. Colored dots indicate phenotype (growth rate and log_10_(*acs*) in **(b)** or log_10_(*ptsG*) in **(c)** of each cell that can grow post-switch. Background shading shows the distribution of phenotypes of all cells in the population.

### Gene regulatory responses are rapid and heterogeneous

Cells varied not only in their ability to grow after the switch, but also in their gene regulatory response: cells near the chamber opening turned on *acs* expression rapidly after the switch, while most cells at intermediate depths did not turn on *acs* expression even after several hours of exposure to acetate (figure 4). One hour post-switch there is a distinct pattern in the chamber: cells near the opening and in the back express *acs*, while cells in the middle do not (figure 4). Considering the GFP maturation time (~20min), cells near the chamber opening turned on *acs* expression within minutes after depletion of glucose, consistent with previous observations in batch culture [39]. Notably, a large fraction of cells that expressed *acs* post-switch could not switch to growth on acetate (figure 2b), consistent with the fact that *acs* expression is a necessary but not a sufficient requirement for growth on acetate [32,39].

**Figure 4.**
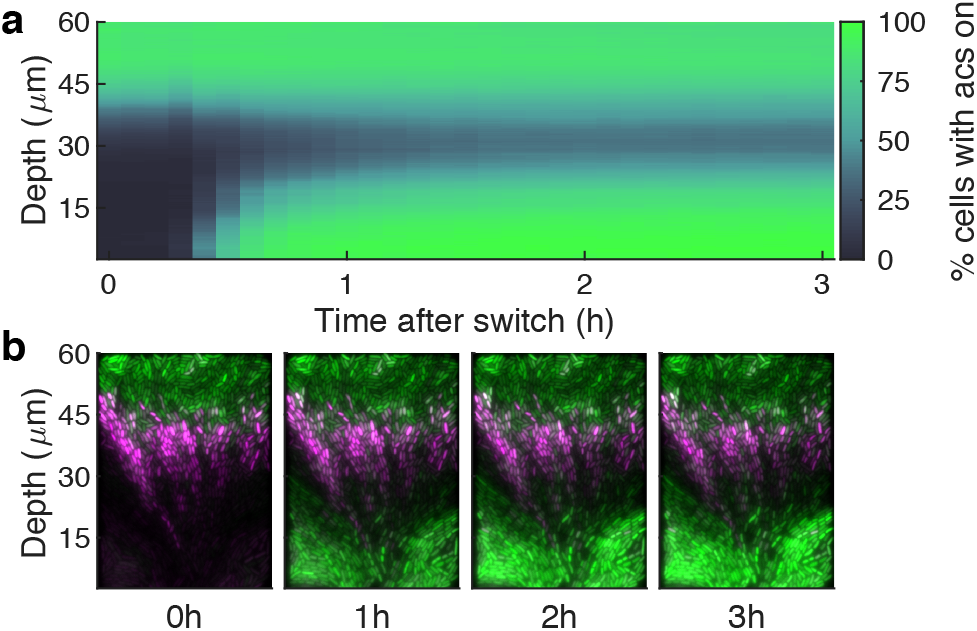
Acs is expressed rapidly after nutrient switch, but not by all cells. **(a)** Cells in the front of the chamber, but not those in the middle, rapidly express *acs* after a switch from glucose to acetate. The percentage of cells expressing *acs* (*acs* >13.5, see Methods) is shown as function of time after the switch and depth. Data from all 15 chambers was pooled together. **(b)** False-color images of a single chamber at four time points; *acs* expression is shown in green, *ptsG* expression in magenta. Brightness and contrast were adjusted for both colors separately, but were not changed between time points.

### Only cells actively growing on acetate can continue growing throughout the nutrient switch

According to our hypothesis, cells that consume acetate pre-switch should not lag. Yet a substantial fraction of cells in the back of the chamber did lag or could not resume growth at all. What distinguishes the few cells that continue growing with no lag? A possible explanation is that these cells are more adapted to growth on acetate before the switch then the rest. Using a logistic regression, we confirmed this hypothesis: cells that continue growing have higher pre-switch growth rates and *acs* expression compared to cells that do lag or that cannot resume growth (SI figure 1, SI table 1,2). Moreover, there is a strong synergistic effect between high growth rates and high *acs* expression: cells that have a high growth rate *and* high *acs* expression have a very high probability to continue growing without lag. Taken together, these finding are consistent with our hypothesis that cells that grow on acetate before the switch can continue growing after the switch, while cells that are not fully adapted to growth on acetate can grow only after a lag phase or not at all.

### Cells growing post-switch have higher *acs* expression and growth rate than their neighbours

Neighboring cells often differ strongly in their ability to grow after the switch to acetate; what is different between the cells that can grow after the switch and their neighbors that cannot? Although neighbouring cells experience approximately the same microenvironment, they can differ in their phenotype because of stochastic gene expression. We thus compared the phenotype of all cells that can grow (with or without lag) after the switch to acetate to the average phenotype of their neighbours (cells within 2μm). Cells that can grow post-switch cluster in two distinct regions of the chamber (back and front of chamber, figure 3a) that differ strongly in their average phenotype. We thus analysed these regions separately, as potentially different factors are important in determining whether a cell can grow after the switch.

We found that cells that could grow post-switch have distinct pre-switch phenotypes compared to their neighbours, but these traits are not the same for the two regions. In the back of the chamber, cells that could grow post-switch had higher *acs* and *ptsG* expression and grew faster than their neighbours (figure 5a). This result is consistent with our hypothesis that exposure to acetate before the switch increases a cell’s chance to grow after the switch. In the front, cells that could grow after the switch have lower *acs* and *ptsG* expression, and higher growth rates than their neighbours (figure 5b). To understand this result, we have to consider that all cells move towards the chamber opening and that, due to differences in friction, the speed of movement varies strongly even between neighbouring cells. As a result, streams of relatively fast-moving cells regularly form [40]. Most cells that can resume growth are located outside of these fast streams, yet their neighbours can be in these streams, and thus have protein expression levels that reflect the microenvironment of regions deeper in the chamber (where *acs* and *ptsG* expression levels are higher).

**Figure 5.**
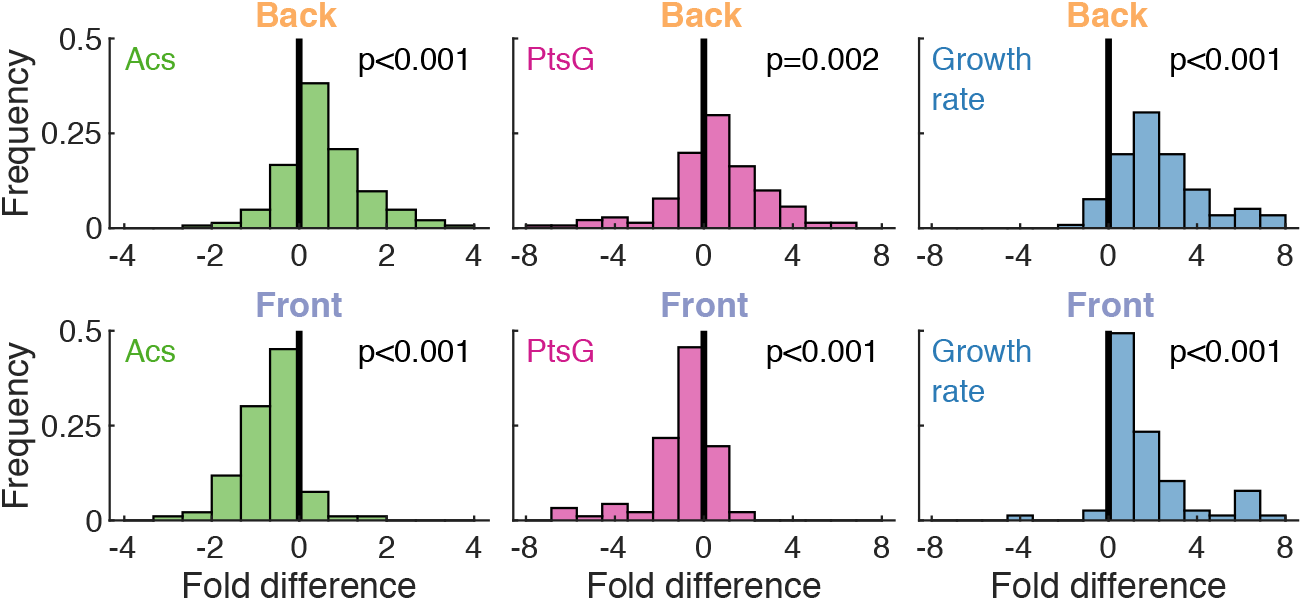
Cells that grow post-switch have higher *acs* expression and growth rate than their neighbors. We compared the phenotype of cells that could grow post-switch with the average phenotype of their neighbors; the analysis was done separately for cells located in the back (top, n=144 cells), or front (bottom, n=94 cells) of the chamber (see figure 3a). The figure shows the fold difference in acs, *ptsG*, and growth rate: negative fold differences indicate that neighbors have on average higher values than the cells that could grow post-switch. p-values show result of two-sided sign test. Cells are considered neighbors if their center-to-center distance is less than 2μm.

### Duration of lag only weakly correlates with pre-switch phenotype

The cells that stopped and resumed growth displayed a wide distribution of lag times (figure 2a). Can we predict how long cells lag from their pre-switch phenotype? To answer this question, we investigated the relation between lag time and pre-switch phenotype (*acs, ptsG*, and growth rate) using a partial correlation analysis. Lag time weakly correlates with *ptsG* expression for cells in the back (with cells expressing more *ptsG* having a longer lag time) and with growth rate for cells in the front (with cells growing faster having a shorter lag time, SI figure 2). However, both correlations are very weak (*ρ*^2^ < 0.11) suggesting that the time required for physiological adaptation is largely independent of the phenotypes we measured.

### Behaviour of distinct subpopulation can explain population response

In summary, the ability of cells to cope with a switch from glucose to acetate depends on their pre-switch *acs* expression and growth rate. Although phenotypes vary continuously between cells, we identified distinct phenotypic classes that can capture most of the variation between cells (figures 2&3). We grouped cells based on their growth rate and *acs* expression and calculate the probability of cells to grow post-switch in each group (*ptsG* expression does not affect these probabilities, SI figure 3). We inferred the most likely metabolic activity of cells in each group based on previous published work (figure 6b).

**Figure 6.**
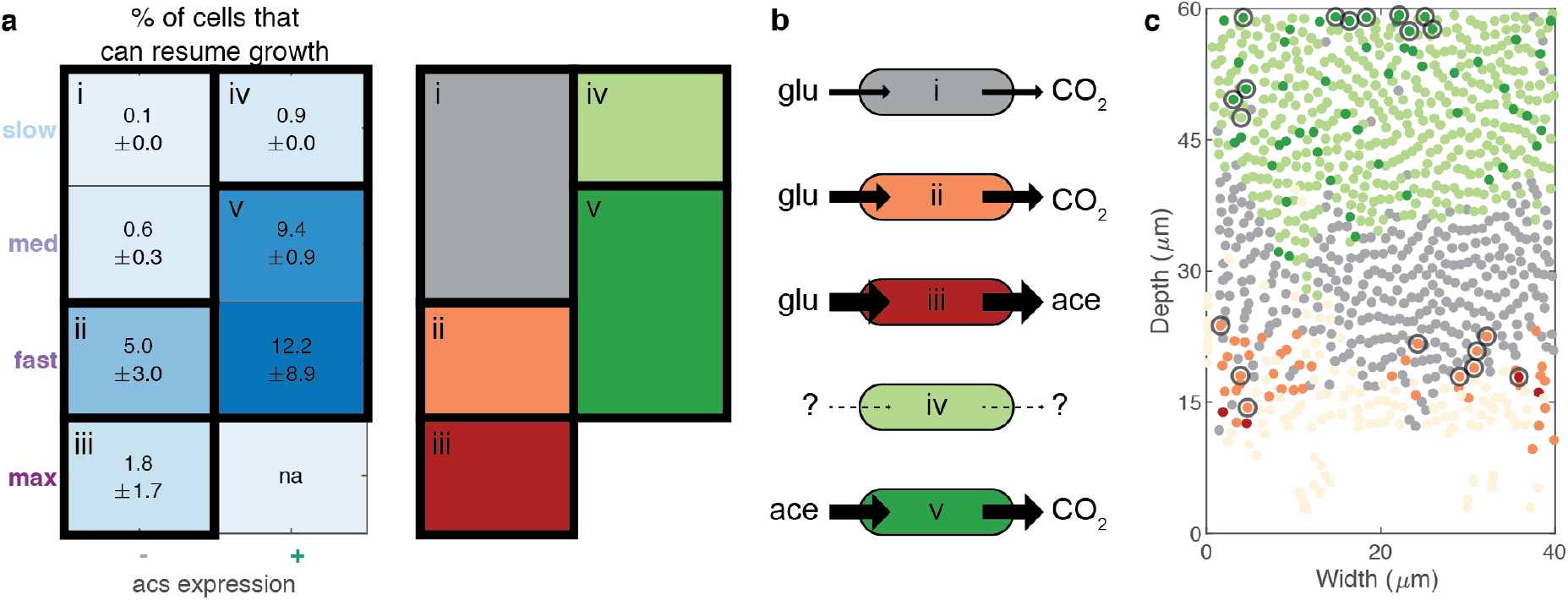
Probability of post-switch growth varies between phenotypic classes. **(a)** We grouped cells based on their *acs* expression and growth rate and calculated for each group the probability for cells to grow post-switch (see Methods). As some cells have unknown growth rates, we estimated the lower and upper bounds of this probability and show the central estimate and range: e.g. a value of 5 ± 3 indicates that the probability to grow post-switch is between 2 and 8%. Five distinct phenotypic classes can capture most of the variation in cells’ phenotype; these classes are indicated by the thick black borders, roman numbers, and schematic on the right. Abbreviations: med: intermediate growth rate; max: maximum growth rate, na: no cells exist in this group. **(b)** Schematic of the inferred metabolic activity of cells in the different classes, based on their observed phenotype and previous published work. Cells in class (i) and (ii) both respire glucose but differ in their growth rate (slow/intermediate and fast, respectively); cells in class (iii) ferment glucose to acetate; cells in class (iv) are likely nutrient starved; and cells in class (v) respire acetate. Thickness of arrows indicates growth rate of cells. **(c)** Location of the different phenotypic classes in space, colors correspond to those in panel b. Pale yellow circles indicate cells with unknown growth rate; most of these are located close to the chamber opening and likely belong to class (ii) or (iii). Cells that could grow post-switch are encircled. The chamber shown is the same as in figure 1a

We found that cells with the highest probability to resume growth were likely growing on acetate before the switch (class v, 9-12%), followed by cells that were growing at high, but non-maximal, rates on glucose (class ii, 5%, figure 6). Cells that were likely starved for carbon (class iv), that were growing slowly on glucose (class i), or fermenting glucose to acetate (class iii), all had very low (<1%) probabilities to grow on acetate after the switch (figure 6). We will now discuss these results in the context of previous studies on the glucose-acetate switch.

We found that most cells expressing *acs* before the switch (class iv & v, figure 6) stopped growing completely after the switch to acetate. *acs* expression is required for catabolism of acetate and energy production, but it is insufficient for anabolic activity (and thus growth), which requires the gluconeogenic pathway and glyoxylate bypass [32,39]. Previous studies have shown that the expression of the glyoxylate pathway can take several hours [39]. Therefore cells that did not express the glyoxylate bypass before the switch, for example because they used small amounts of remaining glucose or other secreted metabolites (e.g. succinate [32]) for anabolic activities, would likely lag for several hours. Previous studies have also shown that cells with insufficient gluconeogenic flux cannot resume growth after a switch to acetate [2]. Therefore, cells that use glucose or are starved for acetate might not be able to resume growth at all after the switch, because they have insufficient gluconeogenic flux. Altogether, results from previous studies might explain why only few cells that express *acs* are able to grow after the switch: these are cells that express the gluconeogenic pathways and the glyoxylate bypass.

We found that, for cells consuming glucose before the switch, the probability for post-switch growth depends on the cell’s growth rate in a non-monotonic way: cells with fast, but not maximal, growth rates (class ii) can grow after the switch to acetate, while very fast (class iii) and very slow (class i) growing cells cannot. How can we explain this non-monotonic dependence of the switching probability on the growth rate? According to previous studies, high growth rates (and high glycolytic activities) imply low gluconeogenic fluxes, thus the fastest growing cells should have the lowest probability to grow on acetate [2,10]. But why can very *slow* growing cells not make the switch? These slow growing cells were under catabolite repression before the switch, as they did not express *acs*, and they likely had low metabolic fluxes and energy reserves. It is thus conceivable that these cells run out of energy before catabolite repression is relieved, which prevents them from switching to growth on acetate. This could also explain why this whole subpopulation could not turn on *acs* expression after the switch (figure 4). Some support for this hypothesis comes from a previous study in *Lactococcus lactis* where it was shown that catabolite repression can prevent cells from switching from growth on glucose to growth on cellobiose [5].

Overall only a fraction of cells in our spatially structured populations could grow after the switch to acetate. For all phenotypic classes this fraction never exceeds 12%, while in a previous study using batch cultures up to 60% of cells could switch from growth on glucose to growth on acetate [2]. This discrepancy might arise from differences between strains or growth media, or might be due to a fundamental difference in the growth environment: spatially structured populations have a hundred to a thousand times higher cell densities than a typical batch culture. In these dense populations, various physical and biochemical interactions can occur between cells that cannot occur in batch cultures [14,15]. These include physical forces between cells [41,42] and secretion of low amounts of metabolites that might affect growth [14,15]. Our study suggests that all these interactions might be essential in determining the response to changing environments of cells living in biofilms and colonies.

## Conclusion

We found that a cell’s ability to cope with a nutrient switch, specifically a switch from glucose to acetate, depends strongly on its pre-switch phenotype. The ability of clonal populations to grow on the new nutrient thus depends on the amount of phenotypic variation within the population. We have shown that phenotypic variation is enhanced in dense spatially structured populations because cells can collectively create heterogeneity in the environment through the uptake and release of compounds. A large heterogeneity in the environment generates a large variety of cell phenotypes that might not occur in batch cultures, but that could be relevant in natural colonies and biofilms. As a result, structured populations might be more able to cope with unexpected environmental fluctuations than homogeneous populations.

We expect our findings to be relevant beyond the glucose-acetate switch in *E. coli:* in nature, many microorganisms live in dense structured populations, where a wide variety of phenotypes are likely to arise. The degree and type of phenotypic variation that arise depend both on environment factors, like the nutrients available, and on biotic factors, like the cellular density and the metabolites excreted. This variation could be advantageous for the population when the environment changes, especially when a subpopulation of cells already expresses the metabolic pathways needed to grow in the new environment. However, if no subpopulations express relevant metabolic pathways before the switch, there could be no advantage of phenotypic variation. Theoretical models could potentially predict the behavior of phenotypically different cells in response to environmental changes. Recently, several models have been developed to study nutrient switches in homogeneous environments [10,43,44] and it would be interesting to extend these models in future work to include the effects of phenotypic variation in the population.

Taken together our results show that it is essential to study microorganisms directly in structured populations to understand how they behave in natural, ever-changing, environments. Our study offers a set of ideas and methods that will help to make progress in this direction. Ultimately, understanding how microorganisms respond to environmental changes will help us to understand and control natural communities and to engineer synthetic communities for specific purposes.

## Methods

### Strains and plasmids

All experiments were done with *E. coli* MG1655 carrying the low copy number pSV66-*acs-gfp-ptsG-rfp* plasmid which contains a GFPmut2 transcriptional reporter for *acs* and a turboRFP transcriptional reporter for *ptsG* [36].

### Media and growth conditions

Cells were grown in M9 medium (47.76 mM Na_2_HPO_4_, 22.04 mM KH_2_PO_4_, 8.56 mM NaCl and 18.69 mM NH_4_Cl) supplemented with 1 mM MgSO_4_, 0.1mM CaCl_2_, 50μg/ml kanamycin (to select for plasmid maintenance), and 0.1% of Tween-20 (Polysorbate-20, to reduce sticking of cells to the sides of microfluidic devices), all from Sigma-Aldrich. This media was supplemented with carbon sources as described below. Overnight cultures were started from a single colony from a LB agar plate and grown in M9 supplemented with 10mM glucose and 5% LB at 37°C in a shaking incubator.

### Microfluidic devices and experiments

The microfluidic devices consist of a series of chambers of 40μm wide, 60μm deep, and 0.76μm high that open on one side into a long (≈2cm) flow channel of 100μm width and 23μm height. The small height of the chambers ensures that cells grow in a monolayer. Molds for the microfluidic devices were fabricated from SU8 photoresist on silicon wafers using a two-layer photolithography process (Microresist, Berlin). Microfluidic devices were manufactured from Polydimethylsiloxane (PDMS, Sylgard 184) as described in reference [36]. The devices were filled with culture medium using a pipette to first wet the channels. An overnight culture of cells was concentrated by centrifugation and loaded into each flow channels by pipetting and cells were pushed into the side chambers. Subsequently the inlets of the flow channels were connected via tubing (Microbore Tygon S54HL, ID 0.76 mm, OD 2.29 mm, Fisher Scientific) to 50ml syringes and media was flown continuously at 0.5ml/h using syringe pumps (NE-300, NewEra Pump Systems).

Cells were first grown in on 10mM glucose until they had filled the chambers (≈18 hours). Subsequently the medium was switched to 800μM glucose for nine hours before switching to 30mM acetate for 38h. Growth media was switched by manual switching tubing between syringes. We calculated that it takes about 18 minutes for the media to change in the flow channel after connecting the tubing to a new syringe and confirmed this by observing how cell length trajectories changed after the switch. We thus set the time of the switch at 18 minutes after the time were we physically changed the tubing.

### Microscopy

Time-lapse microscopy was done using a fully-automated Olympus IX81 inverted microscope, using a 100X NA1.3 oil phase objective (Olympus) and an ORCA-flash 4.0 v2 sCMOS camera (Hamamatsu). For fluorescent images a X-Cite120 120-Watt high pressure metal halide arc lamp (Lumen Dynamics) was used along with the Chroma N49002 (GFP) and N49008 (RFP) filters. Focus was maintained using the Olympus Z-drift compensation system and the microscope was controlled with the CellSens software (Olympus). A microscope incubator (Life imaging services) maintained the sample at 37°C.

### Data acquisition

To measure the pre-switch phenotype, we started imaging two hours before switching to acetate. At this time cells had been exposed to the low glucose medium for seven hours and were fully adapted to the emergent metabolic gradients (in preliminary experiments we established that this process takes about three to four hours). We imaged the population in phase contrast (to measure biomass), GFP (*acs* expression), and RFP (*ptsG* expression) taking an image every six minutes.

### Image analysis

Time-lapse movies where analysed using a combination of Schnitzcell [45], Vanellus (Daan Kiviet, https://github.com/daankiviet/vanellus), Ilastik [46], and customized scripts in Matlab (version 2018b). Movies were registered to compensate for stage movement and cropped to the region of the growth chambers.

### Cell segmentation and tracking

Cells were segmented using supervised machine learning (with Ilastik). Segmentation was done on the superposed phase contrast and fluorescence images. Cell tracking was done using a custom build algorithm that estimates the movement of cells between two subsequent images with optical flow and uses this to predict the location of cells in the subsequent frame [47]. For all cells that could grow after the switch to acetate, we manually corrected the lineage tracking using an adapted version of Schnitzcell [45]. For all other cells, no manual correction was done.

### Gene expression levels

Fluorescent images were deconvolved using the Lucy-Richardson method and background corrected as 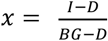, where *I* is the uncorrected intensity, *D* the dark count, and *BG* the background intensity measured in the flow channel. The expression level of *acs* and *ptsG* were estimated for each cell by masking the corrected fluorescent images using the cell-segmentation mask and calculating the average pixel intensity in the GFP and RFP channel, respectively.

### Single cell growth rates

Single cell growth rates (*μ*) are defined as the elongation rate of cells: *l*(*t*) = *l*(0) · *e*^*μ*·*t*^ and estimated as the slope of the linear regression of the log-transformed cell length. The quality of each fit was estimated using the reduced Chi-squared value: 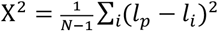, where N is the number of time points over which the regression is done, *l_p_* is the length predicted by the linear regression and *l_i_* is the measured cell length.

We manually corrected the tracking data for all cells that could grow after the switch. For these cells, we measured growth rates by fitting a time window of 5–7 time points (24–36 minutes) ending 6 minutes before the nutrient switch. If a cell was born shortly before the nutrient switch, i.e. if less than 5 time points were available for fitting its growth rate, we used for the first time points the length of the mother cell (*l_mother_*) corrected for asymmetric length division between the focal cell (*l_cell_*) and its sister 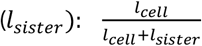, *l_mother_*; we also directly calculated the growth rate of the mother cell just before division, compared this estimate with the one described above and assigned to the focal cell the growth rate with the highest fit quality.

For cells that did not grow after the switch, we automatically corrected tracking mistakes by cutting length trajectories when lengths increased by more than 18% or decreased by more than 15% between two frames (this corresponds to |*μ*| > 1.7 h^−1^). We then performed multiple growth rate fits using time windows of different lengths (N=4–11 frames) at all times between 90 to 0 minutes before the switch. We computed the fits and iteratively compared the quality of the new fit with the best previous candidate; the new fit was considered better if 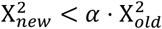, where *α* = {1 if *N_new_* = *N_old_*; 1.25 if *N_new_* > *N_old_*; 0.8 if *N_new_* < *N_old_*}. Visual inspection of these fits suggested that fits over longer time windows were better even if they had a slightly higher X^2^, the factor *α* takes this into account. Finally, if no fit had X^2^ < 10^−3^ we considered the growth rate to be unknown (this threshold was set based on visual inspection). This was the case for 22% of the cells, mostly located close to the chamber opening.

### Identification of cells that could grow after the switch

We identified cells that could grow after the switch to acetate using a combination of automated and manual screening: cells were automatically flagged if their length increased after the switch by 15% relative to the length at the time of the switch, and were subsequently manually screened. Due to imperfections in the automated tracking algorithm, some starting cells could not be identified automatically, therefore we additionally identified all cells that could grow post-switch by visually inspecting the time-lapse movies. Of the 237 cells that could grow after the switch we detected 43% with both methods, 22% only with automatic screening, and 35% only with manual screening.

### Lag time measurement

We considered cells to exit lag phase when they resumed growth, i.e. when they started elongating again. To reduce noise in the growth rate estimates, we first smoothed the log-transformed cell length using a two-hour time window using the *rlowess* (robust local regression) option of the Matlab *smooth* function. Subsequently we estimated growth rates as the slope of the linear regression over a moving widow of five time points (24 minutes), starting at six minutes after the nutrient switch.

The time at which cells exited lag was taken as the first time where their growth rate exceeded 0.02 h^−1^ and remained above this value for at least 75% of the time-points during the next three hours. If a cell exited lag close to its next division (within one hour), we further required that at least one of its daughters’ growth rates was higher than 0.02 h^−1^ directly after birth. If cells divided before exiting lag (n=54), we calculated the lag time of both daughters and assigned the shortest one to the mother cell. Finally, we assigned a lag time of zero to cells that had a growth rate higher than 0.02 h^−1^ at all times. Visual inspection of all length and growth rate trajectories showed that these thresholds could accurately determine the time when cell elongation resumed.

### Discretizing phenotypes into classes

In figures 3 and 5 we grouped cells that could grow post-switch based on their depth in the chamber using k-means clustering with two clusters (threshold depth was 38.5μm). In figure 6 we classified cells based on their phenotype: we discretised gene expression in two classes (on/off) and growth rates in four classes (slow/intermediate/fast/maximum) based on fixed thresholds. We calculated the thresholds for gene expression from the distribution of expression levels of all cells within 15μm of the chamber opening, which generally only express *acs* and *ptsG* at background levels. Cell were considered to have *acs* on if their expression level was three standard deviations above the mean of the background level (*acs*>13.5); cells where considered to have *ptsG* on if their expression level was one standard deviation above the mean (*ptsG*>30.9). The difference between *acs* and *ptsG* reflects the much higher standard deviation in *ptsG* expression levels near the chamber opening. The thresholds for classifying growth rates were calculated using k-means clustering with three clusters, the threshold between *fast* and *maximum* was manually set based on visual inspection. Cells were classified as growing at maximum (*μ* > 0.87 h^−1^), fast (0.40 < *μ* ≤ 0.87 h^−1^), intermediate (0.09 < *μ* ≤ 0.40 h^−1^), or slow (*μ* ≤ 0.09 h^−1^) rate.

### Statistical analysis

We only included chambers that were fully packed with cells during the full observation window, based on visual inspection. Our dataset consists of 15 chambers coming from two independent flow-cells. Given the small number of starting cells per chamber, we analysed cells from all chambers together.

For each phenotypic class, we estimated the probability that cells could grow post-switch. To calculate these probabilities, we considered that for 22% of the cells the growth rate was unknown. We thus calculated an upper and lower bound for these probabilities as 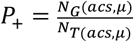 and 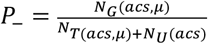, where *N_G_*(*acs,μ*) and *N*_*T*(*acs,μ*)_ are the number of cells that can grow post-switch and the total number of cells with a given *acs* expression and growth rate, and *N_U_*(*acs*) is the number of cell with a given *acs* expression and unknown growth rate. In figure 6 we report the central estimate and range: 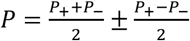.

## Acknowledgements

We thank Daan Kiviet for providing the molds for the microfluidic devices, Jana Fahrion for performing initial exploratory experiments, Alex von Wyl for help with the image analysis, and Susan Schlegel, Glen Dsouza, and the other members from the MSE group for helpful feedback and discussions.

This work was supported by the Swiss National Science foundation (grants no. 31003A_149267 and 31003A_169978 to MA), by an Early Postdoc Mobility fellowship from the Swiss National Science Foundation (grant no. 175123 to SVV), and by ETH Zurich and Eawag.

## Author Contributions

ADC designed the study, collected, analysed, and interpreted the data, and drafted the manuscript; MA designed the study and critically revised the manuscript; SVV designed the study, engineered the genetic constructs, collected, analysed and interpreted the data, and drafted the manuscript. All authors gave final approval for publication and agree to be held accountable for the work performed therein.

## Data Accessibility

The data and code required to reproduce the figures and conclusions are available on the Scholars Portal Dataverse repository: https://doi.org/10.5683/SP2/XVUXDB [48].

