## Supplemental Information for "Metabolic activity affects response of single cells to a nutrient switch in structured populations"

### Supplementary Information

**SI Table 1**. Logistic regression comparing cells that continue growing without interruption (zero lag) after switch from glucose to acetate to all cells in the population. n=10'939; 10'935 degrees of freedom; $\mathrm{X}^2 = 302; p = 4 \cdot 10^{-65}$ versus constant. See also SI figure 1a.

| | *Estimate* | *Standard error* | *t-Statistic* | *p-Value* |
|---|---|---|---|---|
| *Intercept* | -7.3 | 0.33 | 22 | $1 \cdot 10^{-105}$ |
| *Growth rate* | 2.9 | 0.71 | 4.0 | $6 \cdot 10^{-5}$ |
| *acs* | 0.031 | 0.0051 | 6.0 | $2 \cdot 10^{-9}$ |
| *Groth rate:acs* | 0.11 | 0.022 | 5.3 | $1 \cdot 10^{-7}$ |

**SI Table 2**. Logistic regression comparing cells that continue growth without interruption (zero lag) after switch from glucose to acetate to all cells that have finite, but non-zero, lag. n=235; 231 degrees of freedom; $\mathrm{X}^2 = 112;\ p = 4 \cdot 10^{-24}$ versus constant. See also SI figure 1b.

| | *Estimate* | *Standard error* | *t-Statistic* | *p-Value* |
|---|---|---|---|---|
| *Intercept* | -2.8 | 0.73 | -3.9 | $1 \cdot 10^{-4}$ |
| *Growth rate* | -0.94 | 1.4 | -0.67 | 0.5 |
| *acs* | 0.0035 | 0.012 | 0.30 | 0.8 |
| *Groth rate:acs* | 0.25 | 0.053 | 4.6 | $4 \cdot 10^{-6}$ |

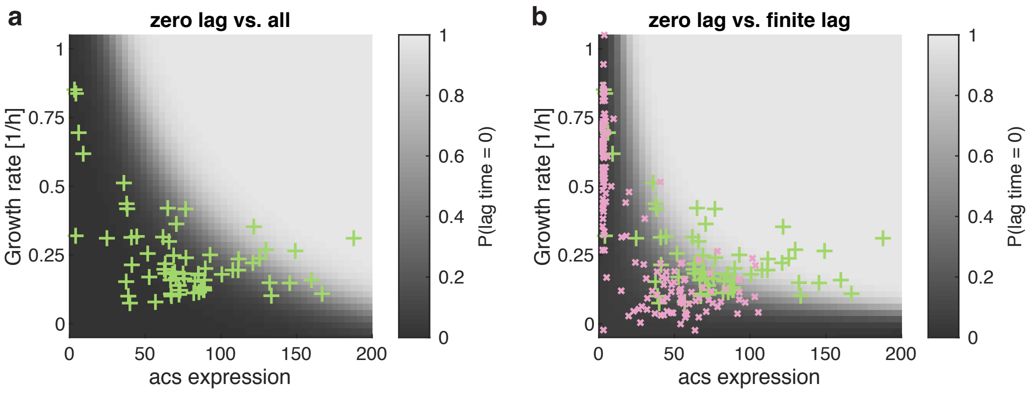


**SI Figure 1. Cells with zero lag have high *acs* expression and growth rate. (a):** Phenotype (*acs* expression and growth rate) of cells that continue growing without interruption (zero lag) after switch from glucose to acetate (green +). Background shading shows probability ($\log_{10}$ transformed) that a cell has zero lag. This probability was calculated from a logistic regression comparing cells with zero lag to all cells in the population (see SI Table 1). Higher *acs* expression and higher growth rates synergistically increase the probability that a cell will have zero lag. **(b):** Same as in a, but comparing cells that have zero lag (green +) to those that have non-zero, but finite lag (magenta x). This probability was calculated from a logistic regression comparing cells with zero lag to all cells with non-zero, but finite lag.

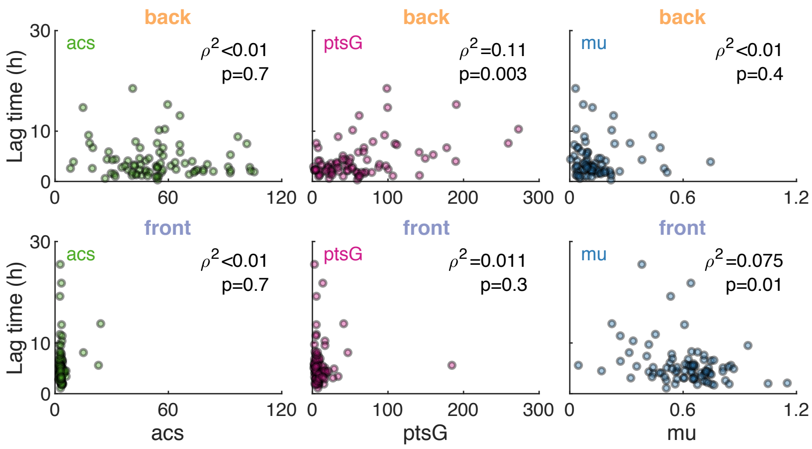


**SI Figure 2. Pre-switch phenotype weakly predicts duration of lag.** For cells which resume growth after a lag, the lag time was compared to the pre-switch phenotype (*acs*, *ptsG*, and growth rate). The analysis was done separately for cells in the back (top, n=80 cells) and front (bottom, n=88 cells) of the chamber (see figure 3a). A partial correlation analysis shows that in the back, lag time is weakly correlated with *ptsG* expression, while in the front the lag time is weakly anti-correlated with growth rate. $\rho^2$ and p values of Spearman rank-based partial correlation are shown.

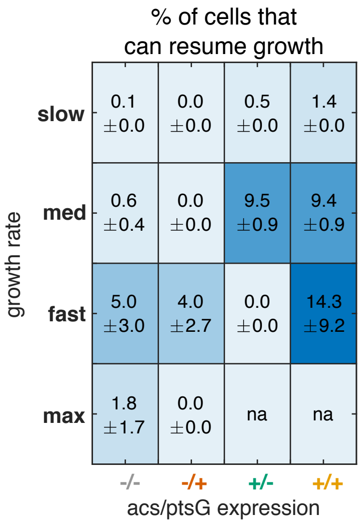


**SI Figure 3. *ptsG* expression does not affect probability of post-switch growth.** Same as figure 6a, but including *ptsG* expression in the classification of cells into discrete phenotypic groups. *ptsG* generally does not affect the probability of post-switch growth. The only exception is fast growing cells that express *acs*, however in this group only 8 cells were able to grow post-switch (all expressing *ptsG*). Values show the probability that cells in a given phenotypic group can grow post-switch, in the format central estimate $\pm$ range (e.g. $5 \pm 3$ indicates probability between 2 and 8%). Abbreviations: med: intermediate growth rate, max: maximum growth rate, and na: no cells exist in this group.

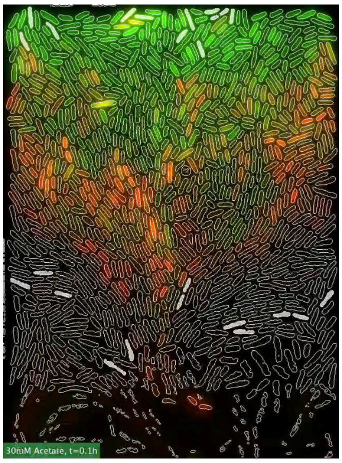


**SI Movies 1-4. Cells response during glucose-acetate switch in structured population.** The movies show the response of cells in a chamber during a switch from glucose to acetate; the time and growth medium is shown in bottom left corner. The expression of *acs* (green) and *ptsG* (magenta) are shown, brightness and contrast are adjusted at each frame separately. Cell outlines (from cell segmentation) are shown as white lines. We successfully identified all cells that could restart growing after the switch and most of their offspring (partly translucent-white mask).
